# Sensory progenitors influence patterning of the mammalian auditory sensory epithelium

**DOI:** 10.1101/2023.11.13.566920

**Authors:** Caryl A. Young, Emily Burt, Vidhya Munnamalai

## Abstract

During embryonic development Wnt signaling has been shown to influence proliferation and sensory formation in the cochlea. How the dual nature of Wnt signaling is coordinated is unknown. In this study, we define a novel role for a Wnt regulated gene, *Mybl2,* which was already known to be important for proliferation, in influencing patterning and determining the size of the sensory epithelium in the murine cochlea. Using a quantitative spatial analysis approach and analyzing *Mybl2* loss-of-function cochleas, we show that *Mybl2* simultaneously specifies the progenitor niche and the size of the sensory domain, and influences the positioning of the medial sensory domain boundary via *Jag1* regulation during the mid-gestational stages. *Mybl2* conditional knockout resulted in a decrease of proliferation within the progenitor niche. During the late embryonic stages, conditional knockout of *Mybl2* produced a wider sensory epithelium across the radial axis with an increase in ectopic inner hair cell formation. These data suggest that *Mybl2*-positive progenitors play a role in boundary formation and patterning the sensory epithelium.

**Summary Statement:** *Mybl2* is a Wnt-regulated gene encoding a transcription factor that is expressed in the cochlear progenitor niche and influences the boundary formation between the niche and the sensory domain during mid-cochlear developmental stages, thereby impacting the size of the sensory epithelium.

## 1. Introduction

The sensory epithelium of the cochlea, the organ of Corti, has a highly specialized organization consisting of one row of sound-detecting inner hair cells and three rows of sound-amplifying outer hair cells across its radial axis. This organization of the sensory epithelium confers auditory processing by guiding afferent and efferent innervation to target inner hair cells and outer hair cells. The sensory epithelium is flanked by the inner sulcus (IS) epithelium along the medial edge of the coiled cochlea and the outer sulcus (OS) epithelium on the lateral edge of the coiled organ. Proper formation of the sensory epithelium and its organization is dependent on the correct placement of the sensory epithelial boundaries. However, the molecular mechanisms that determine their formation and their boundaries are unclear. Several studies suggest that multiple signaling pathways influence the formation of the medial versus the lateral compartments of the cochlea (Basch et al., 2016; Ellis et al., 2019; Hayashi et al., 2007; Huh et al., 2012; Hwang et al., 2010; Jansson et al., 2019; Munnamalai and Fekete, 2016; Munnamalai and Fekete, 2020; Munnamalai et al., 2012; Ohyama et al., 2010; Oliver et al., 2021; Petrovic et al., 2015).

Our previous studies showed that during development, the Wnt signaling pathway influenced the formation of the medial compartment, which comprises the IS domain and the medial sensory domain (MS) in the mouse cochlea or specified neural cell fates in the chicken cochlea (Munnamalai and Fekete, 2016; Munnamalai et al., 2017; Oliver et al., 2021). We also showed that the Wnt secretion enzyme, PORCN was enriched on the medial half of the cochlea between embryonic day (E)13.5-E14.5 (Oliver et al., 2021) during the stages when the IS, MS and lateral sensory (LS) and OS domains are established; thus, WNT ligand secretion is optimally positioned to promote the formation of the medial IS and MS domains, especially because WNT morphogens are classically considered to be short range morphogens (Routledge and Scholpp, 2019). For example, in the intestine, WNT ligands are secreted from the progenitor niche and are only dispersed via the action of cell division (Farin et al., 2016). As in several developing systems, activation of the Wnt pathway stimulated cell cycle re-entry in the cochlea (Jacques et al., 2012; Liu et al., 2022a; Samarajeewa et al., 2019; Samarajeewa et al., 2018). Several studies showed that the Wnt pathway also regulates the expression of *Jag1,* or *Serrate1* expression in the mouse and chicken cochleas respectively, which suggests that there is conservation in the regulation of the sensory domain by the Wnt pathway (Jacques et al., 2012; Munnamalai and Fekete, 2016; Munnamalai et al., 2017). However, it is unknown how the Wnt pathway can confer both progenitor and sensory specification.

Our recent studies showed that the SOX2-positive sensory domain undergoes refinement to be repositioned from the medial edge on E12.5 to the center of the cochlear duct floor by E14.5 (Thompson et al., 2023). The SOX2-positive sensory domain will continue to form the sensory epithelium. Upon differentiation, the support cells retain SOX2 expression, while the hair cells do not (Dabdoub et al., 2008). In the E14.5 mouse cochlea, the medial IS domain houses a niche of progenitor cells adjacent to the sensory domain. In this study, we investigate potential molecular mechanisms via an uncharacterized gene in the cochlea, *Mybl2,* by which the Wnt pathway simultaneously regulates the proliferation in the IS domain and the size of the JAG1-mediated SOX2-positive sensory domain in the developing mouse cochlea. During the early otic stages, the Wnt pathway positions the neurosensory domain in the inner ear (Zak and Daudet, 2021). Similarly, we determine whether Wnt-regulated MYBL2 plays a novel role in influencing the positioning of the boundary between the IS domain and the sensory domain on E14.5.

## 2. Results

To determine how the Wnt pathway influences boundary formation between the IS domain and the cochlear sensory epithelium, we first compared the spatial position of the prosensory domain on E12.5 relative to E14.5 (Figure 1). We immunolabeled for JAG1 and SOX2 expression, two sensory domain markers, on E12.5 and E14.5 cochleas (Figure 1A-1D). On E12.5, JAG1 expression was positioned on the medial edge of the cochlea (Figure 1A). Since JAG1-Notch signaling regulates the expression of SOX2 and the formation of sensory epithelia (Kiernan et al., 2001; Kiernan et al., 2006), the SOX2-positive sensory domain was also located on the medial edge of the cochlea (Figure 1B). By E14.5, both JAG1 and SOX2 expressions were refined to re-position the sensory domain at the center of the cochlear duct floor (Figure 1C, 1D). How this refinement and the repositioning of the JAG1-positive and SOX2-positive sensory domain occurs is unknown. To determine the spatial relationship between Wnt signaling via PORCN expression and the Wnt-regulated JAG1 expression on E14.5, we performed spatial analysis on PORCN by co-immunolabeling E14.5 cochlear sections for PORCN, JAG1, and E-cadherin. E-cadherin was utilized as a marker for the LS domain (Chacon-Heszele et al., 2012; Whitlon, 1993). We found that PORCN expression was enriched in the medial half of the cochlea, and JAG1 expression lies within the PORCN expression domain (Figure 1E). However, there was a large medial region where PORCN expression is present, but JAG1 expression was absent (Figure 1E, arrow). As previously shown, E-cadherin was enriched in the LS (Figure 1F). At this stage, JAG1 was absent in the lateral domains of the cochlear epithelium on E14.5; thus, JAG1 expression showed no overlap with E-cadherin. We performed quantitative spatial analysis of PORCN, JAG1, and E-cadherin by immunofluorescence (Figure 1G), which suggested that although JAG1 expression lies within the PORCN domain, there must be some unknown factor that suppressed JAG1 expression in the medial IS domain. Quantitative spatial analysis also showed that E-cadherin peak expression lies in the LS domain where neither PORCN nor JAG1 are expressed (Figure 1G). *Jag1* was found to be a direct Wnt target gene in hair follicles (Estrach et al., 2006) and several studies showed that *Jag1* expression was positively regulated by Wnt activation in the E12.5 cochlea (Jacques et al., 2012; Munnamalai and Fekete, 2016). We found Wnt regulation of *Jag1* on E12.5 was consistent when we analyzed JAG1 expression in E12.5 *Emx2Cre; β−catenin* (*β−cat*) conditional knockout (cKO) cochleas (Figure S1) as *Emx2* is expressed in the early cochlea by E12.5 (Holley et al., 2010) (Figure S1A). In control cochleas, JAG1 is expressed on the medial side, but is downregulated in *β-cat* cKOs (Figure S1B, 1C). JAG1 fluorescence was decreased by 71% ± 8.8% (mean ± SEM, p-value = 1.04e-7) in the *β-cat* cKOs relative to littermate control cochleas (Figure S1D). However, by E14.5, JAG1 was no longer expressed at the medial edge, despite the expression of PORCN (Figure 1E, 1G), which suggests that PORCN/ WNT regulation of JAG1 expression was dynamic (Munnamalai and Fekete, 2013; Munnamalai and Fekete, 2016; Oliver et al., 2021). This observation was consistent with our previous studies which showed that *Jag1* responsivity to Wnt activation was different at different stages of cochlear development (Munnamalai and Fekete, 2016). A 6-hour bath treatment of E12.5 cochleas *in vitro* by CHIR99021, a GSK-3β inhibitor that activates the Wnt pathway, showed an up regulation of *Jag1*. However, when explants were treated one day later *Jag1* expression was suppressed.

**Figure 1:**
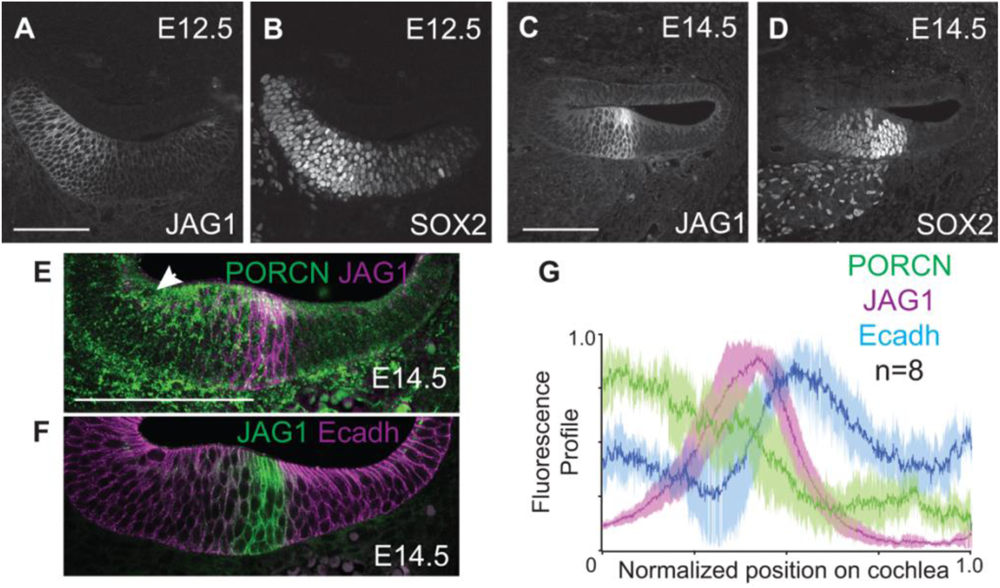
The sensory domain is refined between E12.5 and E14.5. (A, B) JAG1 and its downstream effector, SOX2 were both enriched at the medial edge of the cochlear duct on E12.5. (C, D) By E14.5, the JAG1 and SOX2 domains were repositioned to the middle of the duct. (E) PORCN was enriched in the medial half of the cochlea. JAG1 was expressed within the PORCN domain on E14.5. (F) JAG1 and E-cadherin have complementary, non-overlapping expression patterns on E14.5. (G) Spatial profile analysis of PORCN, E-cadherin, and JAG1 expression on E14.5 revealed their relative expression patterns. (N = 8 cochleas). Scale bar = 100μm

To determine whether JAG1 was dynamically regulated by the Wnt pathway in a spatiotemporal manner, we used *Isl1Cre* to induce later *β-cat* cKO as *Isl1Cre; Tdt* showed strong expression in the mid-turn by E14.5 (Figure S2), where we performed quantitative analysis of domain sizes (Figure 2). We analyzed JAG1 expression in *Isl1Cre*; *β-cat* cKO cochleas on E14.5, at which timepoint, the JAG1-positive and SOX2-positive domains were refined away from the medial edge of the cochlea (Figure 1C, 1D). In the E14.5 control cochlea, the JAG1 domain, which labels the MS domain, was positioned in the center of the cochlear duct floor (Figure 2A). As previously shown, E-cadherin expression was absent in the JAG1-positive MS domain but was enriched in the LS domain (Figure 2A’). In E14.5 *Isl1Cre*; *β-cat* cKO cochleas (Figure 2B), JAG1 expression was not reduced as we saw in E12.5 *Emx2Cre; β-cat* cKO cochleas (Figure S1). We quantified the temporal impact of *β-cat* cKO between E13.5-E14.5 using *Isl1Cre* by measuring the widths of the radial domains in the E14.5 cochlea. To determine how each domain is influenced by temporal *β-cat* cKO on E13.5, we measured the total width of the cochlea, the widths of the medial and lateral compartments, and the widths of each individual domain; the IS, MS, LS and OS (Figure 2C). The MS domain width was measured using JAG1 as a marker and the IS domain width was measured from the medial boundary of the MS domain to the luminal medial edge of the cochlea. The width of the LS domain was measured using E-cadherin and SOX2, as shown in Figure 2C, and the OS domain width was measured from the lateral SOX2 boundary to the luminal lateral edge of the cochlea. The total radial width of E14.5 control cochleas were measured to be 192.24 μM ± 3.12 μM, while the total radial width of E14.5 *Isl1Cre*; *β-cat* cKO cochleas were significantly decreased to 177.91 μM ± 3.42 μM (p-value = 4.88e-5) (Figure 2D). However, the medial and lateral compartments themselves did not significantly change in the *Isl1Cre*; *β-cat* cKO cochleas relative to control cochleas (Figure 2E, 2F). The widths of the IS and MS domain were measured and normalized to the width of the medial compartment (IS + MS domains). The IS domain was larger and occupied 71% ± 3% of the medial compartment, while the MS domain was smaller and occupied 29% ± 3% of the medial compartment in control cochleas (Figure 2G, 2H). In the *Isl1Cre*; *β-cat* cKO cochleas, the IS domain decreased in size to occupy 52% ± 4% of the medial compartment, while the MS domain expanded to occupy 48% ± 4% of the medial compartment (p-value = 2.02e-6) (Figure 2G, 2H). Thus, there was a significant increase in the size of the JAG1-positive MS domain in *Isl1Cre*; *β-cat* cKO cochleas relative to controls. This was also evident from the increase in the region where E-cadherin expression was absent in the MS domain (Figure 2B’). The widths of the LS and OS domains were not significantly different between control and *Isl1Cre*; *β-cat* cKO cochleas (Figure 2I, 2J). The increase in the size of the MS domain comes at the expense of the size of the IS domain on E14.5. This led us to hypothesize that there exists a Wnt-regulated gene that is expressed in the IS domain on E14.5 that suppresses the expression of JAG1 to drive the refinement of JAG1 from the medial edge of E12.5 to the center of the cochlear duct floor on E14.5 (Figure 1A-1D).

**Figure 2:**
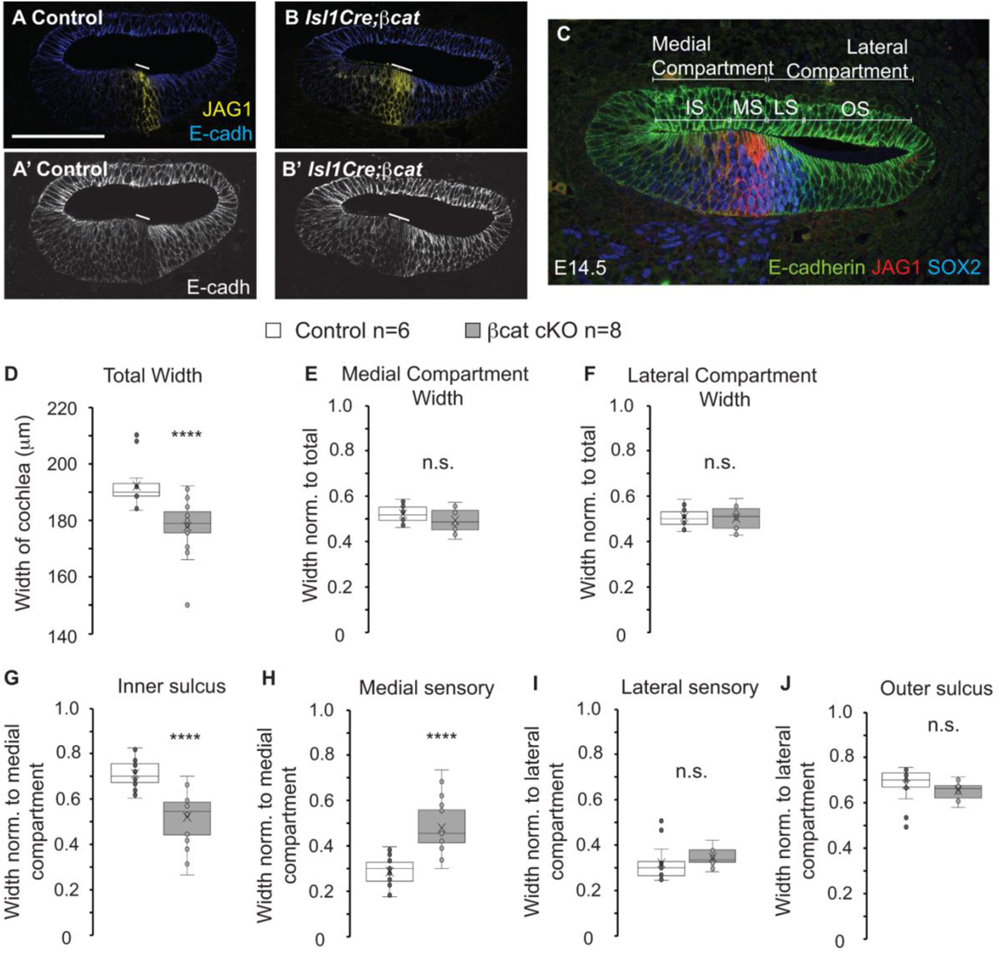
JAG1 is spatiotemporally regulated by Wnt signaling. (A) Immunolabeling for JAG1 and E-cadherin in E14.5 control cochlea. (A’) E-cadherin was absent in the MS domain, enriched in the LS domain and was complementary. (B) JAG1 expression in *Isl1cre*; *β-cat* cKO cochlea on E14.5 was not reduced but expanded. (B’) E-cadherin was further downregulated in the region of JAG1 expansion. (C) Indication of domains and markers used. JAG1 labels the MS domain and defines the IS-MS boundary. E-cadherin is enriched in the LS domain. SOX2 is enriched in the sensory domain (MS + LS). The lateral boundary of SOX2 and E-cadherin segregate the LS and OS domains. (D-J) Quantification of domain sizes in control and *Isl1cre; β-cat* cKOs on E14.5. (Control N = 6, n = 15 sections, *β-cat* cKO N = 8, n = 19 sections) A two-tailed students t-test was performed for each quantification and Bon Ferroni correction factor was applied for multiple comparisons for determining significance. (D) Total Width p-value = 4.88e-5 (E) Medial Compartment Width p-value = 0.08, (F) Lateral Compartment Width p-value = 0.89, (G) Inner Sulcus p-value = 2.02e-6, (H) Medial Sensory p-value = 2.02e-6, (I) Lateral Sensory p-value = 0.21, (J) Outer Sulcus p-value = 0.21. Solid line in A, A’, B, B’ marked the MS domain. P-value < 1e-4 indicated by ****. Scale bar = 100μm

To identify a potential Wnt-regulated gene, we performed RNA-sequencing on E14.0 cochleas that were treated with CHIR99021 for 6 hours. We isolated genes that encode for transcription factors with a log-fold change greater than 0.5 and examined their spatial expression by *in situ* hybridization in E14.5 cochlear tissue. Consistent with PORCN expression on E14.5, *Mybl2* encoded a transcription factor that was found to be expressed on the medial side of the cochlea, where PORCN expression was the highest. MYBL2 is most known for its role in regulating the cell cycle during development and disease (Liu et al., 2022b; Musa et al., 2017; Papetti and Augenlicht, 2011; Ward et al., 2018).

To determine whether the Wnt pathway spatially regulates *Mybl2* expression, we performed quantitative spatial analysis using immunofluorescence staining for PORCN and *in-situ* hybridization of *Mybl2* on E14.5 wildtype cochleas (Figure 3). PORCN was restricted to the medial half of the cochlea on E14.5 (Figure 3A). We then performed *in situ* hybridization for *Mybl2* and found that *Mybl2* is enriched on the medial side of the cochlear duct floor (Figure 3B). We compared the spatial profiles of both PORCN and *Mybl2* and found that *Mybl2* expression lies within the PORCN expression domain (Figure 3C). These data support that Wnt spatially regulates *Mybl2* expression in the medial side of the cochlea during development. To verify that *Mybl2* was regulated by the Wnt pathway, we generated E14.5 *Isl1Cre*; *β-cat* cKO embryos to examine *Mybl2* expression by *in situ* hybridization in the cochlea (Figure 3D, 3E). *Mybl2* expression remained in the medial side of control cochleas (Figure 3D, dotted lines and arrow), while in *Isl1Cre*; *β-cat* cKO cochleas, *Mybl2* expression was absent (Figure 3E, arrow). Therefore, the Wnt pathway promotes the expression of *Mybl2* in the medial side of the cochlea.

**Figure 3:**
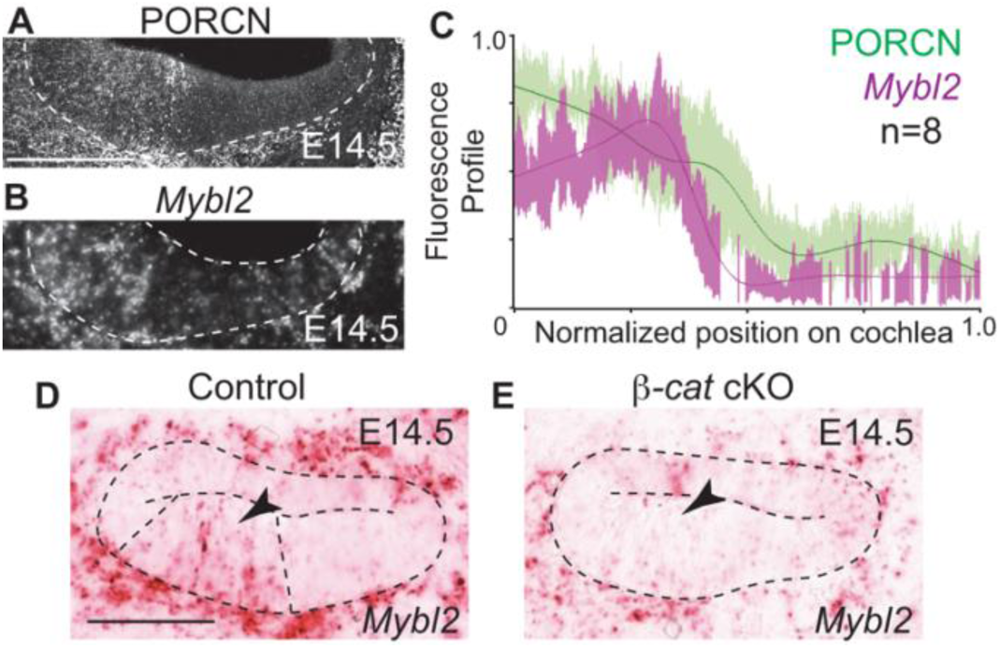
The Wnt pathway regulates *Mybl2* in the cochlea. (A) Immunolabeling of PORCN on E14.5 was enriched on the medial half of a wildtype cochlea. (B) *Mybl2* transcript expression in an E14.5 wildtype cochlea. (C) Quantitative spatial profile comparing PORCN and *Mybl2* expression on E14.5 (N= 8 cochleas). (D-E) *Mybl2* expression in control E14.5 cochlea and in its littermate *Is1lCre;β-cat* cKO cochlea on E14.5. Scale bar = 100μm

The medial, spatial expression of *Mybl2* made it a suitable candidate as a Wnt-regulated gene encoding a transcription factor that could suppress JAG1 expression on the medial side of the cochlea. We performed quantitative spatial analysis of JAG1 (Figure 4A) relative to *Mybl2* expression (Figure 4B) in the E14.5 wildtype cochlea. A single trace of JAG1 and *Mybl2* (Figure 4A, 4B) showed that they were not expressed in the same domains of the cochlea. The lateral boundary of *Mybl2* expression abutted the medial boundary of the JAG1-positive MS domain (Figure 4C). *Mybl2* expression is enriched in the medial domain but there was a sharp decline in *Mybl2* expression (Figure 4C, arrowhead) where JAG1 expression peaks within the MS domain; thus, based on its expression relative to JAG1, *Mybl2* expression was located in the IS domain. This spatial expression across the radial axis remained consistent in profile plots averaged across 8 cochleas such that *Mybl2* expression was restricted to the IS domain, while JAG1 marked the MS domain (Figure 4D). Thus, *Mybl2* and JAG1 form a boundary to give rise to the IS and MS domains respectively.

**Figure 4:**
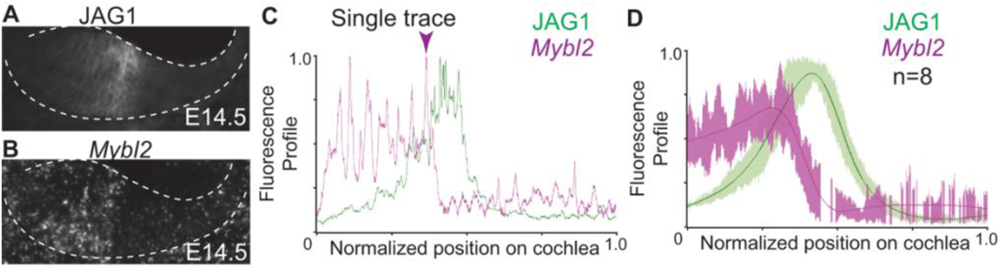
Spatial juxtaposition of JAG1 and *Mybl2* in the E14.5 cochlea. (A) Immunolabeling for JAG1 on E14.5 exhibited its central positing on the cochlear duct floor. (B) *Mybl2* transcripts on E14.5 were present medially. (C) Representative single quantitative spatial profile plot between JAG1 and *Mybl2*. (D) Average quantitative spatial profile plot between JAG1 and *Mybl2* (N = 8 cochleas). Scale bar = 100μm

Since MYBL2 is appropriately positioned in the IS domain to potentially reposition JAG1 expression to the center of the cochlear duct on E14.5, we examined the size of the JAG1-positive MS domain in *Mybl2* cKO cochleas. To test this, we generated E15.5 Sox2*Cre^ER^*; *Mybl2* cKO embryos that were induced daily with tamoxifen between E10.5 to E14.5. Embryos were harvested on E15.5 and cochleas were immunolabeled for JAG1 and SOX2 to measure domain sizes (Figure 5), as previously shown in E14.5 *Isl1Cre; β-cat* cKO cochleas (Figure 2). In E15.5 control cochleas, JAG1 expression was refined to the center of the cochlear duct floor (Figure 5A). In *Sox2Cre^ER^; Mybl2* cKO cochleas, the JAG1 domain was expanded (Figure 5B). Since SOX2 is downstream of JAG1 and specifies the sensory domain, we immunolabeled for SOX2 in control and *Mybl2* cKO cochleas (Fig. 5C, D). There were no significant differences between E15.5 control and *Sox2Cre^ER^; Mybl2* cKO cochleas in the total radial widths of the cochlea (Figure 5E), the widths of the medial compartment (Figure 5F), or the widths of the lateral compartment (Figure 5G). When we measured the widths of the IS and MS domains normalized to the total width of the medial compartment, we found that the IS domain decreased in size from occupying 57% ± 3% of the medial compartment in control cochleas to occupying 38% ± 4% of the medial compartment in *Sox2Cre^ER^; Mybl2* cKO cochleas (Figure 5H). On the other hand, the MS domain was expanded from occupying 43% ± 3% in the control cochleas to 62% ± 4% in *Mybl2* cKOs (p-value = 1.28e-10) (Figure 5I). This result followed the same trends that we observed in the E14.5 *Isl1Cre; β-cat* cKO cochleas (Figure 2). The widths of the LS and OS domains were not significantly different compared to control cochleas (Figure 5J, 5K). Lastly, we measured the total SOX2 area in control and *Sox2Cre^ER^; Mybl2* cKO cochleas and we found a significant increase in SOX2 area by 18% ± 8% (p-value = 1.25e-4) in the *Mybl2* cKO cochleas relative to controls (Figure 5L). Similar results were found for both domain sizes and SOX2 area using different Cre-expressing mouse lines (data not shown). These data showed that *Mybl2* plays a role in establishing the size of the prosensory domain during development.

**Figure 5:**
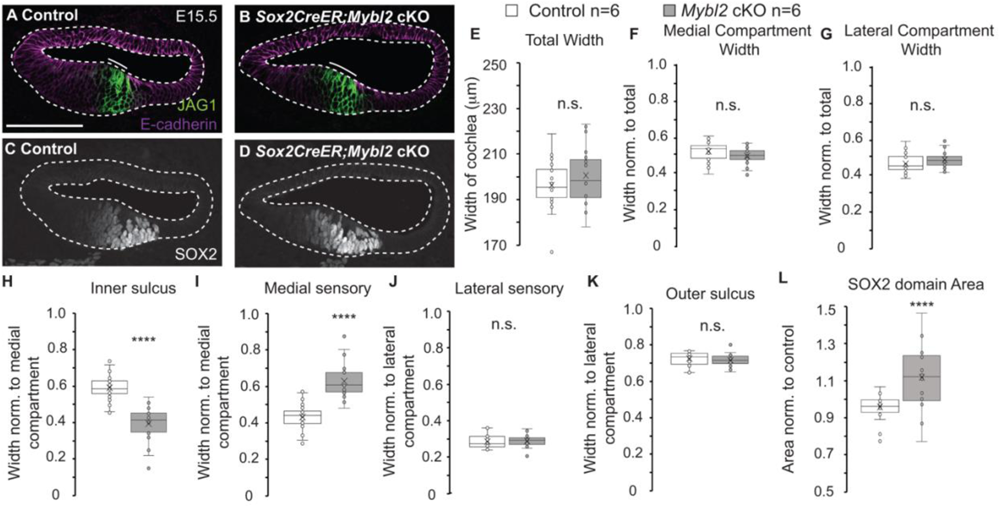
*Mybl2* specifies the size of the medial prosensory domain. (A) Central positioning of JAG1 domain in the E15.5 control cochlea. (B) E15.5 *Sox2Cre^ER^; Mybl2* cKO cochleas exhibited an expanded JAG1 domain. (C) SOX2 immunolabeling in control cochlea on E15.5. (D) SOX2 was expanded in *Sox2Cre^ER^; Mybl2* cKO cochleas. (E-L) Quantification of domain sizes in control and *Sox2Cre^ER^; Mybl2* cKOs on E15.5 (Control N = 6, n = 24 sections, *Mybl2* cKO N = 6, n = 24 sections). A two-tailed students t-test was performed for each quantification and Bon Ferroni correction factor was applied for multiple comparisons for determining significance. (E) Total Width was not significant, p-value = 0.17 (F-G) Medial Compartment Width, p-value = 0.12 and Lateral Compartment Width, p-value = 0.12 were not significant. (H) Inner Sulcus p-value = 1.28e-10, (I) Medial Sensory domain p-value = 1.28e-10. (J) Lateral Sensory domain p-value = 0.53. (K) Outer Sulcus p-value = 0.53, (L) SOX2 domain p-value = 1.25e-4. P-value < 5e-4 indicated by ****. Scale bar = 100μm

The role of MYBL2 was already known to be essential for proliferation and cell cycle progression (Musa et al., 2017; Papetti and Augenlicht, 2011). To rule out that the expansion of the SOX2 domain was not due to an increase in proliferation by other means independent of MYBL2, we investigated the impact of the loss of *Mybl2* on proliferation in the cochlea (Figure 6). Ki67, a marker for cells in mitosis, was expressed in cells of the IS domain in control cochleas (Figure 6A), in the same domain where *Mybl2* was expressed (Figure 6B). We generated E15.5 *Sox2CreER; Mybl2* cKO embryos and analyzed changes in proliferation by Ki67 expression in E15.5 cochleas (Figure 6C, 6D). On E15.5, there was a 22% ± 6% (p-value = 9.33e-6) decrease in Ki67 labeling in the IS domain of *Mybl2* cKO cochleas, suggesting a decrease in proliferation upon the loss of *Mybl2*.

**Figure 6:**
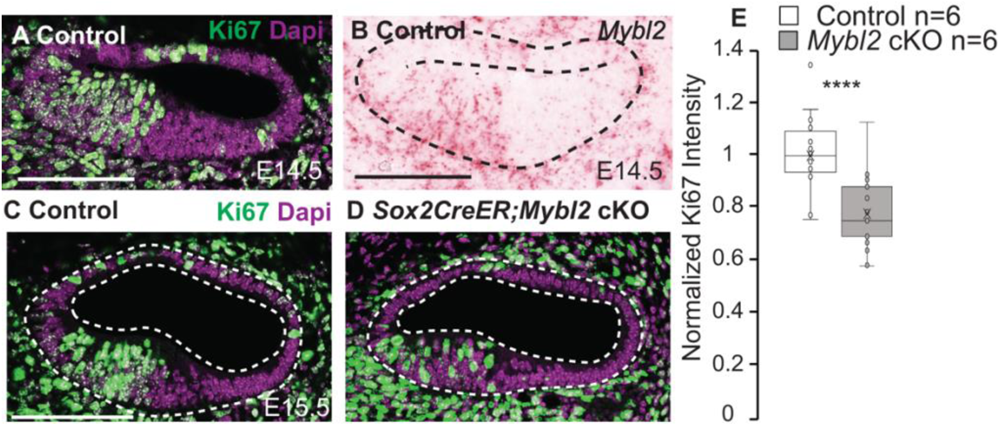
*Mybl2* is expressed and regulates proliferation in the progenitor niche in the E14.5 cochlea. (A) Ki67, a mitotic marker immunolabels proliferating cells in the E14.5 cochlea. (B) *Mybl2* transcripts were expressed in the same region as the progenitor niche. (C) Littermate control cochlea immunolabeled for Ki67 on E15.5. (D) Ki67 labeling was decreased in *Sox2Cre^ER^; Mybl2* cKO cochleas. (E) Quantification of total fluorescent Ki67 levels in control and *Sox2Cre^ER^; Mybl2* cKO cochleas on E15.5 (Control N = 6, n = 22 sections, *Mybl2* cKO N = 6, n = 21 sections). A two-tailed students t-test was performed, and Bon Ferroni correction factor was applied for multiple comparisons for determining significance. p-value = 9.33e-6. P-value < 5e-4 indicated by ****. Scale bar = 100μm

To determine whether the JAG1 and SOX2 boundary effects leading to an expanded sensory domain that we observed on E15.5 influenced the final outcome on cochlear patterning, we analyzed E18.5 *Sox2Cre^ER^*; *Mybl2* cKO cochleas (Figure 7). Tamoxifen was administered once daily from E10.5 until E14.5 and harvested on E18.5. On E15.0, *Mybl2* was no longer expressed in the cochlea; therefore, tamoxifen induction was not required after E14.5. We immunolabeled whole mount cochleas for Myosin VI (MYO6) and SOX2 to label the hair cells and sensory epithelium, or support cells respectively. Upon comparing the littermate control and E18.5 *Sox2Cre^ER^; Mybl2* cKO cochleas, we observed an expanded SOX2 domain and an overall increase in SOX2 intensity (Figure 7A-7D). In addition to the expanded sensory epithelium, we also observed ectopic inner hair cells resulting in the form of hair cell doublets in *Mybl2* cKO cochleas (Figure 7C, arrows and 7C’). For more precise analysis of the effects on the SOX2-positive epithelium, we generated optical sections of the z-stacks of whole mount cochleas to quantify the changes in SOX2 area and SOX2 intensity on these (Figure 7A’-7D’) because we observed an accumulation of SOX2-positive cells medially in the *Mybl2* cKO cochleas, which would contribute to changes in area (Figure 7D’, magenta arrow). Both area and intensity of SOX2 differed in the control cochleas based on their longitudinal location in the base, mid and apex regions (data not shown). Thus, we quantified SOX2 in the basal, mid, and apical regions (Figure 7E, 7F) and we found a significant increase in SOX2 area and SOX2 intensity in all regions of the cochlea, with the largest increase observed in the base and smallest increase observed in the apex of the *Mybl2* cKO cochleas compared to the controls. In the base we measured a 47% ± 16% (p-value = 7.8e-9) increase in SOX2 area, and a 59% ± 21% (p-value = 2.93e-8) increase in SOX2 intensity. In the mid-region of the cochlea, the loss of *Mybl2* resulted in a 33% ± 12% (p-value = 3.58e-7) increase in SOX2 area, and a 43% ± 18% (p-value = 3.19e-6) increase in SOX2 intensity. Lastly, in the apex, we measured a 20% ± 11% (p-value = 1.44e-4) increase in SOX2 area and a 23% ±16% (p-value = 1.8e-3) increase in SOX2 intensity in *Mybl2* cKOs compared to their control littermates. Although the average increase was the smallest in the apex, we did observe maximum increases of 50% in SOX2 area and 72% in SOX2 intensity in the apex within individual sections. Thus, the loss of *Mybl2* in the apex is capable of producing larger areas of sensory epithelia. Despite previous reports that the *Sox2Cre^ER^* mice results in *Sox2* haploinsufficiency and their cochleas showed a decrease in *Sox2* levels relative to their ‘no Cre’ littermates (Atkinson et al., 2018); we observed a significant increase in SOX2 in *Sox2Cre^ER^; Mybl2* cKO cochleas, which suggests that there was positive feed-forward loop to compensate and further enhance SOX2 in *Sox2Cre^ER^; Mybl2* cKO cochleas beyond the ‘no Cre’ control cochleas.

**Figure 7:**
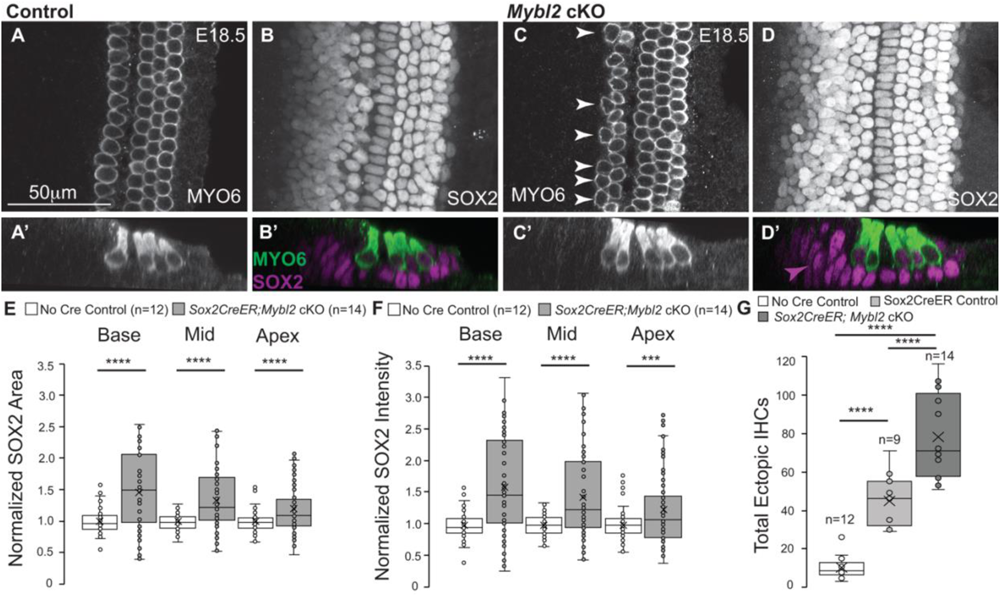
*Mybl2* is important for mature patterning of the sensory epithelium. A) MYO6 immunolabels the single row of inner hair cells and three rows of outer hair cells in whole mount control cochlea on E18.5. (A’) Optical cross-section of a control cochlea. (B) SOX2 immunolabels the sensory epithelium, or the support cells in whole mount control cochlea on E18.5. (B’) Optical cross-section of MYO6 and SOX2 labeling. (C) *Sox2Cre^ER^; Mybl2* cKOs on E18.5 show ectopic inner hair cells in an additional row, labeled by MYO6. (C’) Optical cross-section of C. (D) SOX2 staining in *Sox2Cre^ER^; Mybl2* cKOs reveals an expanded sensory domain with increased SOX2 levels on E18.5. (D’) Optical cross-section of D with MYO6 and SOX2 immunolabeling. Arrowhead shows medial expansion of SOX2-positive cells. (E-F) Quantification of SOX2 area and SOX2 intensity in control and *Sox2Cre^ER^; Mybl2* cKO cochleas on E18.5 in base, mid, and apex. (Control N = 12 cochleas, *Mybl2* cKO N = 14 cochleas, base n = 69 optical sections, mid n = 63 optical sections, apex n = 72 optical sections). Two-tailed Students t-test performed for statistical significance. (E) SOX2 area quantification (base p-value = 7.8e-9, mid p-value = 3.58e-7, apex p-value = 1.44e-4) (F) SOX2 intensity quantification (base p-value = 2.93e-8, mid p-value = 3.19e-6, apex p-value = 1.83e-3. (G) Quantification of total ectopic inner hair cells in ‘no Cre’ control cochlea, *Sox2Cre^ER(+/-)^*control cochlea, and *Sox2Cre^ER^; Mybl2* cKO cochleas. Tukeys test performed comparing *Sox2Cre^ER^; Mybl2* cKO (p-value = 6e-11) and *Sox2Cre^ER(+/-)^* control (p-value = 2.04e-4) to ‘no Cre’ control cochlea and *Sox2Cre^ER^; Mybl2* cKO (p-value = 2.1e-4) to *Sox2Cre^ER(+/-)^* control. (‘no Cre’ Control N = 12 cochleas, *Mybl2* cKO N = 14 cochleas, *Sox2Cre^ER(+/-)^*control N = 9 cochleas. Bon Ferroni correction factor was applied for multiple comparisons for determining significance. P-value < 0.001 indicated by ***, p-value < 5e-4 indicated by ****. Scale bar = 50μm

Although the primary effect of the loss of *Mybl2* was on the SOX2-positive sensory epithelium, we also quantified the number of inner hair cell doublets in control and *Mybl2* cKO cochleas (Figure 7G). We harvested E18.5 cochleas from *Sox2Cre^ER^; Mybl2* cKO embryos and their ‘no Cre’ control littermates, and separately generated E18.5 cochleas from *Sox2Cre^ER(+/-)^* heterozygous ‘control’ embryos. We included *Sox2Cre^ER(+/-)^* cochleas as the appropriate control since previous reports showed the formation of ectopic inner hair cells due to *Sox2* haploinsufficiency (Atkinson et al., 2018) because excess SOX2 itself is prohibitive to hair cell formation (Dabdoub et al., 2008). Comparing the cochleas from these three groups and performing a one-way ANOVA, we found that inner hair cell doublets increased to 78.5 ± 6.3 in *Sox2Cre^ER^; Mybl2* cKO cochleas (p-value = 6e-11, Tukey test) and inner hair cell doublets increased to 45 ± 4.9 in *Sox2Cre^ER^*^(+/-)^ control cochleas (p-value = 2.04e-4, Tukey test) compared to the ‘no Cre’ control cochleas that had 10.5 ± 1.8 inner hair cell doublets (Figure 7G). The increase in inner hair cell doublets in *Sox2Cre^ER^; Mybl2* cKO cochleas were significantly higher than the inner hair cells doublets in the *Sox2Cre^ER^*^(+/-)^ control cochleas (p = 2.1e-4, Tukey test).

Upon measuring the total length of the cochleas, the *Mybl2* cKOs were significantly shorter by 15% compared to the ‘no Cre’ littermate controls (p-value = 6e-7, Tukey post-hoc test), but were only 8% significantly shorter compared to the *Sox2Cre^ER(+/-)^* control cochleas (p-value = 0.005, Tukey post-hoc test). We then counted the number of aligned inner hair cells and the total number of inner hair cells (aligned + ectopic) in the ‘no Cre’ control, *Sox2Cre^ER^*^(+/-)^ control and the *Sox2Cre^ER^; Mybl2* cKO cochleas. There were no significant differences in the number of aligned inner hair cells per 1000 μm between the three groups. If the cochlea length was decreased compared to the *Sox2Cre^ER^*^(+/-)^ control cochleas but the number of aligned inner hair cells do not change, then we would expect to see a decrease in the total number of inner hair cells in *Sox2Cre^ER^; Mybl2* cKO cochleas relative to *Sox2Cre^ER^* control cochleas. However, we did not observe any significant change in the total number of inner hair cells per 1000 μm, which suggests that the number of ectopic inner hair cells that were formed in the *Sox2Cre^ER^; Mybl2* cochlea was indeed higher than in the *Sox2Cre^ER^*^(+/-)^ control. In fact, the number of ectopic inner hair cells that were formed in *Mybl2* cKOs increased by 74% relative to *Sox2Cre^ER^*^(+/-)^ control cochleas, despite the shortened cochlea by 8%.

## 3. Discussion

The Wnt pathway is most known for its proliferative role during development and disease (Chai et al., 2012; Chai et al., 2011; Jacques et al., 2012; Munnamalai and Fekete, 2013; Munnamalai and Fekete, 2016; Ng et al., 2019). In both the mouse and chicken inner ears, pharmacological activation of the Wnt pathway and overexpression of WNT ligands induced proliferation (Jacques et al., 2012; Munnamalai et al., 2017; Stevens et al., 2003). As a result, the Wnt pathway holds promise for promoting regeneration in the mammalian cochlea to restore hearing (Jan et al., 2013; McLean et al., 2017; Samarajeewa et al., 2019; Samarajeewa et al., 2018). However, recent studies in the cochlea showed that the Wnt pathway is also necessary for early sensory specification through the regulation of *Jag1* expression. JAG1-Notch signaling then regulates the expression of SOX2 to establish the sensory domain and eventually, the sensory epithelium. Other studies showed that the Wnt pathway also promoted differentiation (Davidson et al., 2012; Shi et al., 2010; Shi et al., 2014) and the Wnt pathway likely possesse stage-dependent roles (Munnamalai and Fekete, 2013). Thus, it is important to understand the multifaceted roles underlying Wnt signaling. *Jag1* has been shown to be a direct Wnt target gene (Estrach et al., 2006). Consistent with this, our previous studies and several others also showed that treatment of mouse cochleas with a Wnt activator, or infection of chicken cochleas with a *Wnt-ligand* expressing virus showed an increase in *Jag1*, or the chicken homologue, *Serrate1* expression (Jacques et al., 2012; Munnamalai and Fekete, 2016; Munnamalai et al., 2017). Both are genes expressed on the neural side of the cochlea (also known as the medial side in the mouse cochlea). Our previous studies showed that although there are several WNT ligands expressed in the cochlea, the WNT secretion enzyme (Proffitt and Virshup, 2012), PORCN is spatially enriched on the medial side, which suggests there is higher WNT secretion on the medial side (Oliver et al., 2021). Thus, WNT secretion would be higher on the medial side to specify both proliferation and sensory specification. However, we showed that on E14.5, the proliferating progenitor pool is segregated from the JAG1-positive MS domain. This suggests that proliferation and sensory specification are two distinct processes that are co-regulated by separate Wnt-mediated mechanisms. Given that WNT morphogens are short-range signaling molecules (Farin et al., 2016; Routledge and Scholpp, 2019), we examined the spatial context mid-development on E14.5 of PORCN relative to both *Mybl2*-positive progenitors and the JAG1-positive MS domain in the mouse cochlea. Both the progenitor niche and JAG1 expression lie in the PORCN expressing domain on the medial half of the mouse cochlea and are within positional range to be regulated by the Wnt pathway.

Previous studies quantitatively showed that the SOX2 domain that was once located on the medial edge on E12.5 was gradually repositioned to the center of the cochlear duct floor by E14.5 (Thompson et al., 2023). We showed that it is during this stage that a prominent pool of progenitors occupy the IS domain where PORCN expression is enriched and is adjacent to the SOX2-positive cells in the sensory domain (Oliver et al., 2021). This led us to postulate that Wnt signals within the progenitors influenced the boundary between the IS domain and the sensory domain to segregate the progenitor niche from the sensory domain. Thus, we sought to identify the molecular mechanism by which the Wnt pathway can regulate two distinct processes simultaneously while also refining the JAG1 domain and establishing the IS-sensory domain boundary. Consistent with previous studies that showed early Wnt activation led to an increase in JAG1 expression (Jacques et al., 2012; Munnamalai et al., 2017; Zak and Daudet, 2021), we showed that early deletion of *β-catenin* resulted in a loss of JAG1 expression (Figure S1). However, we also showed that JAG1 is dynamically regulated by the Wnt pathway, which suggested there was a Wnt-regulated suppressor of *Jag1* expression. Consistent with this, a later temporal conditional deletion of β-catenin from E13.5 did not abolish JAG1 expression, but instead led to an expansion in the size of the JAG1-postive MS domain at the expense of the IS domain and an increase in the area of the SOX2-positive sensory domain. These data suggest that there is an unknown Wnt-regulated gene expressed in the IS domain on E13.5-E14.5 that refines and re-positions the JAG1-positive MS domain. Our Wnt-transcriptomic analysis and *in situ* hybridization studies revealed that *Mybl2*, a previously uncharacterized gene in the cochlea, is expressed in the IS domain. *Mybl2*, which encodes a transcription factor, is also known for its role in regulating cell cycle progression.

Based on quantitative spatial analysis of *Mybl2* relative to JAG1, we hypothesized that MYBL2 was expressed in the right location to influence the formation of the medial boundary of the MS domain, while maintaining a pool of proliferating cells in the IS domain. This would imply that, apart from its role as a cell cycle regulator, MYBL2 plays a novel function in sensory epithelial patterning by influencing the positioning of the sensory medial boundary. In support of this hypothesis, MYBL2 ENCODE ChIP-seq data showed *Jag1* to be an MYBL2 target gene. When we analyzed *Mybl2* cKO cochleas on E15.5, we found that there was an increase in the area of the SOX2-positive sensory domain. It is interesting to note that this increase in SOX2 area occurs despite a decrease in Ki67-positive proliferating cells in the IS domain in *Mybl2* cKO cochleas. The SOX2 domain extends into the IS domain due to a relief in JAG1 suppression and a medial shift in the IS-MS domain boundary. In this way, the Wnt pathway can simultaneously regulate a progenitor niche by regulating *Mybl2* expression, while regulating *Jag1* expression in the MS domain. However, it is through MYBL2, that the Wnt pathway determines the size of the MS domain (Figure 8). Interestingly, we found no significant quantitative changes in the lateral domains of the cochlea with the loss of *Mybl2*.

**Figure 8:**
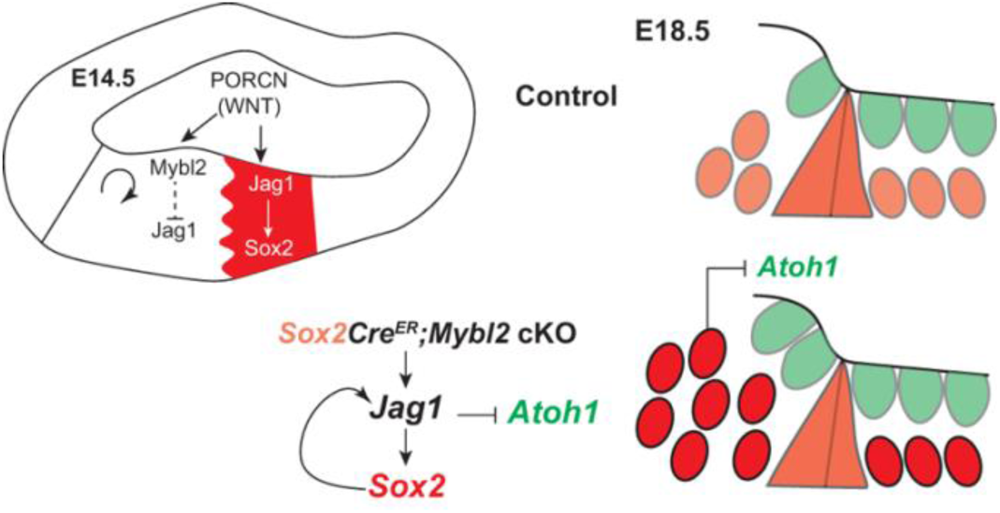
Model for Wnt mediating dual roles in proliferation and patterning via regulation of *Mybl2* in the cochlea. Wnt influences proliferation and patterning of the medial compartment via regulation of *Mybl2*. MYBL2 inhibits JAG1 expression to specify the IS and MS domains. *Mybl2* cKO results in increase in the size of the sensory domain.

We expected that the increase in SOX2 area on E15.5 would leave a lasting impact on E18.5. As such when we analyzed E18.5 *Sox2CreER; Mybl2* cKO cochleas, we found that both the size of the SOX2-positive sensory epithelium and SOX2 levels were significantly enhanced compared to ‘no Cre’ littermate control cochleas. These effects on the sensory epithelium were consistent from the basal, mid, and apical regions of the cochlea. Previous studies showed that *Sox2* haploinsufficiency was caused by the *Sox2Cre^ER(+/-)^* line resulting in a decrease in *Sox2* expression compared to ‘no Cre’ littermate controls (Atkinson et al., 2018). However, in our experimental paradigm, because the conditional deletion of *Mybl2* expanded the SOX2 domain (using two different Cre lines), this created a feed-forward loop to enhance *Sox2*-mediated Cre recombination, ultimately resulting in compensation and an increase of SOX2 above that of levels seen in the ‘no Cre’ control cochleas (Figure 8).

As a consequence of the expanded sensory epithelium and a shift in the medial boundary of the MS domain, we expected that there could be an accompanying increase in the inner hair cells. Excess SOX2 can also be repressive to *Atoh1* expression (Dabdoub et al., 2008) and it is likely that the *Sox2* haploinsufficiency and the reported decreases in SOX2 levels in *Sox2Cre^ER(+/-)^* cochleas was permissive to ectopic inner hair cell formation (Atkinson et al., 2018). However, in the context of *Sox2Cre^ER^; Mybl2* cKO cochleas, we saw a significant increase in levels and expansion of the SOX2 sensory epithelium on E18.5. We found a significant increase in the formation of ectopic hair cells, but these effects were not due to haploinsufficiency, but due to the enhanced SOX2 levels. In fact, we surmise that the *Sox2Cre^ER^; Mybl2* cKO cochleas have an even greater potential for the formation of inner hair cells if it were not for the possible repression of *Atoh1* expression by excess SOX2 (Figure 8). Despite this increase in SOX2, we still saw a 74% greater increase of ectopic inner hair cells as a downstream effect of expanding the sensory epithelium in *Sox2Cre^ER^; Mybl2* cKO cochleas relative to *Sox2Cre^ER+/-^* cochleas.

In conclusion, the Wnt pathway is indeed capable of simultaneously regulating two different processes: proliferation and sensory domain specification in a spatial manner and establishing the boundary in between through the regulation and action of MYBL2. Since both are regulated by the Wnt pathway, and one domain feeds forward to suppress another, this resembles subdomain formation by an incoherent feed-forward loop. We also identify a novel role of MYBL2 as a transcription factor that mediates boundary formation and patterning of the sensory epithelium spatially by suppressing JAG1 expression. Thus, the cochlear sensory progenitors play an unexpected role in influencing patterning of the mammalian auditory sensory epithelium. One possibility for the need for such a role by progenitors is to maintain their ‘stem cell-like’ characteristics to maintain developmental plasticity and replace damaged or dying cells during the developmental process until the sensory epithelium has fully formed.

## 4. Materials and Methods

### Animal Husbandry

*β−catenin flox* (B6(Cg)-Ctnnb1^tm1Knw^/J, Jax Strain #:022775) and *Mybl2 flox (*B6;B6CB-Mybl2, RIKEN, Japan) mice were used to generate cKOs. *β−cat* mice were crossed with *Emx2Cre* (B6.Cg-Emx2<tm2(cre)Sia>/SiaRbrc, RIKEN, Japan) mice for early cochlear cKO by E12.5, and *Isl1Cre* (Isl1^tm1(cre)Sev^/J, JAX Strain #:024242) for later cochlear knockout in the mid-turn by E13.5. *Emx2Cre*; *β−cat* cKO embryos were harvested on E12.5. *Isl1Cre; β−cat* cKO embryos were harvested on E14.5, late in the day. *Mybl2 flox* mice were crossed with *Sox2Cre^ER^*(B6;129S-^Sox2tm1(cre/ERT2)^Hoch/J JAX Strain #:017593) to generate cKOs. Mice were timed-mated, and the day a plug was observed was designated E0.5. Pregnant dams were administered tamoxifen (1.5mg/ 10g body weight) once daily between E10.5-E14.5 to generate *Sox2Cre^ER^; Mybl2* cKOs. *Sox2Cre^ER^; Mybl2* cKO embryos were harvested on E15.5 and E18.5. Swiss Webster wildtype mice (Envigo, Indianapolis, IN, USA) were used to generate E14.5 embryos for expression studies. Ai9 (B6.Cg-Gt(ROSA)26Sor^tm9(CAG-tdTomato)Hze^/J, Jax Strain #: 007909) mice were used to show Tdt expression upon Cre recombination. Sequencing data at these stages did not reveal sex differences; therefore, we indiscriminately pooled data from all embryos. All animal procedures were performed in accordance with the Institutional Animal Care and Use Committee (IACUC) guidelines at The Jackson Laboratory.

### Histology

Embryos were decapitated and heads were fixed in 4% paraformaldehyde (PFA) (Electron Microscopy Sciences, Hatfield, PA, USA) overnight at 4°C. Embryo heads were processed through 10%, 20%, and 30% sucrose solutions and cryofrozen in tissue freezing medium (TFM) (General Data Healthcare, Cincinnati, OH, USA). Cochleas of E18.5 embryos were fixed in 4% PFA and dissected as wholemounts.

### Immunofluorescence and in situ hybridization

Sectioned tissues were blocked with 2% donkey serum (Jackson Immuno-Research, West Grove, Pennsylvania, USA) in PBS/0.5% Triton for 1 hour at room temperature followed by primary antibody incubation overnight at 4°C. The following day, tissues were incubated with Alexa-conjugated secondary antibodies for 2 hours at room temperature (Invitrogen) and nuclei were counterstained with DAPI (Abcam, Waltham, MA, USA). Tissues were mounted with Fluoromount G mounting medium (Life Technologies Corporation, Carlsbad, CA, USA). Primary antibodies included: rat anti-E-cadherin (1:250, AB11512, Lot# 1012145-3, Abcam, Waltham, MA, USA), goat anti-Jagged1 (1:250, sc-6011, Lot# I2115 Santa Cruz Biotechnology, CA, USA), rat anti-Ki67 (1:250, Ref:14-5698-82, Lot# 2196796 Invitrogen, Waltham, MA, USA), goat-anti SOX2 (1:500, AF2018, Lot# KOY0421101, R& D Systems, Minneapolis, MN, USA), rabbit anti-SOX2 (1:500, ab97959, Lot# GR3244869-1,Abcam, Cambridge, UK), rabbit anti-Myosin VI (1:500, Ref: 25-6791, Proteus, Waltham, MA, USA), rabbit anti-PORCN (1:250, PA5-43423, Invitrogen, Waltham, MA, USA). Antigen retrieval was performed for Ki67 immunolabeling with 10mM sodium citrate/ 0.05% Tween 20 buffer at pH 6.0 for 15 minutes at 99°C. RNAscope for *Mybl2* was performed according to the manufacturer’s protocol (Advanced Cell Diagnostics, Newark, CA, USA). Probe used to detect transcripts: Mm_Mybl2 Ref # 563211.

### Data Acquisition and Image processing

Image acquisition was performed on a Zeiss LSM800 confocal microscope (ZEISS, Oberkochen, BW, Germany) at The Jackson Laboratory Microscopy core and on a brightfield/ epifluorescence Olympus BX51 microscope with a Spot insight CMOS camera were used to acquire images at 20X and 40X. Z-stacks were analyzed using the FIJI software (NIH). Measurements were made on raw image data. For figure preparation, TIFF images were processed in Adobe Photoshop and assembled in Adobe Illustrator (Adobe, San Jose, CA, USA).

### Data Analysis and Quantification

Spatial profile analyses were performed along the floor of the cochlear duct using the FIJI software. Profile expression data were plotted along the total widths of the respective cochleas, averaged across samples, and plotted in R. The total width of the cochlea was measured from the medial edge to the lateral edge on the luminal side. Fluorescent intensity data was normalized between 0 and 1 with intensity minima at 0 and intensity maxima at 1. Profile plots were compiled across 8 cochleas.

Domain size measurements were performed using FIJI. Thresholding was used to determine comparable fluorescent levels demarcating domain boundaries and the widths of the domains were measured between these boundaries. The widths of the IS and MS domains were normalized to the width of the medial compartment of the cochlea and the widths of the OS and LS domains were normalized to the width of the lateral compartment of the cochlea. JAG1 immunolabeled the MS domain, E-cadherin and SOX2 immunolabeling were used to determine the lateral boundary of the LS. The size of the total SOX2 domain and total Ki67 fluorescent levels on E15.5 was determined in FIJI by setting a minimum threshold and measuring the area and intensity of the signal respectively. On E18.5, orthogonal projections were generated in FIJI, a threshold was set, and the total SOX2 area and intensity was measured. All area and total intensity data were normalized to the respective control mean values. The ectopic inner hair cells are hair cells that were observed medially to the main, aligned, single row of inner hair cells and were counted along the total length of the cochlea. Whole mount images were shown as both maximum projections and orthogonal optical sections.

Quantifications were performed on a minimum of 15 sections from at least six cochleas per condition (exact number of sections and cochleas are stated in the legends of each figure). All data fit normal distributions. Statistical analysis between control and cKO cochleas was performed in R by independent, two-tailed Student’s *t* test. To compare the three groups on E18.5 (‘no Cre’ control, *Sox2Cre^ER^*^(+/-)^, and *Sox2Cre^ER^; Mybl2* cKOs) we performed a One-way ANOVA, followed by a post-hoc, Tukey test to compare each group to one another. Corrected p-values were calculated and applied for multiple comparisons.

## 5. Acknowledgements

The authors acknowledge the Microscopy Core at The Jackson Laboratory for equipment maintenance, upkeep, and assistance. The authors thank members of the Munnamalai lab for providing critical feedback on the manuscript.

## 7. Author Contributions

C.A.Y., study conceptualization and design, development of methodology, acquisition, data analysis, writing and revision of the manuscript. E.B., study design, acquisition, data analysis, and revision of the manuscript. V.M., funding, material support, study conceptualization and design, development of methodology, acquisition, analysis, and interpretation of data; writing and revision of the manuscript. All authors read and approved the final manuscript.

## 8. Ethics Approval

All experiments were performed under IACUC protocol 18013: PI Vidhya Munnamalai, in compliance with the U.S. Department of Health and Human Services and reviewed by The Jackson Laboratory Institutional Animal Care and Use Committee.

## 9. Data Availability

The datasets used and/ or analyzed during the current study are available from the corresponding author on request.

## References

1. Atkinson, P. J., Dong, Y., Gu, S., Liu, W., Najarro, E. H., Udagawa, T. and Cheng, A. G. (2018). Sox2 haploinsufficiency primes regeneration and Wnt responsiveness in the mouse cochlea. J Clin Invest 128, 1641–1656.

2. Basch, M. L., Brown, R. M., 2nd, Jen, H. I., Semerci, F., Depreux, F., Edlund, R. K., Zhang, H., Norton, C. R., Gridley, T., Cole, S. E., et al. (2016). Fine-tuning of Notch signaling sets the boundary of the organ of Corti and establishes sensory cell fates. Elife 5.

3. Chacon-Heszele, M. F., Ren, D., Reynolds, A. B., Chi, F. and Chen, P. (2012). Regulation of cochlear convergent extension by the vertebrate planar cell polarity pathway is dependent on p120-catenin. Development 139, 968–978.

4. Chai, R., Kuo, B., Wang, T., Liaw, E. J., Xia, A., Jan, T. A., Liu, Z., Taketo, M. M., Oghalai, J. S., Nusse, R., et al. (2012). Wnt signaling induces proliferation of sensory precursors in the postnatal mouse cochlea. Proc Natl Acad Sci U S A 109, 8167–8172.

5. Chai, R., Xia, A., Wang, T., Jan, T. A., Hayashi, T., Bermingham-McDonogh, O. and Cheng, A. G. (2011). Dynamic expression of Lgr5, a Wnt target gene, in the developing and mature mouse cochlea. J Assoc Res Otolaryngol 12, 455–469.

6. Dabdoub, A., Puligilla, C., Jones, J. M., Fritzsch, B., Cheah, K. S., Pevny, L. H. and Kelley, M. W. (2008). Sox2 signaling in prosensory domain specification and subsequent hair cell differentiation in the developing cochlea. Proc Natl Acad Sci U S A 105, 18396–18401.

7. Davidson, K. C., Adams, A. M., Goodson, J. M., McDonald, C. E., Potter, J. C., Berndt, J. D., Biechele, T. L., Taylor, R. J. and Moon, R. T. (2012). Wnt/beta-catenin signaling promotes differentiation, not self-renewal, of human embryonic stem cells and is repressed by Oct4. Proc Natl Acad Sci U S A 109, 4485–4490.

8. Ellis, K., Driver, E. C., Okano, T., Lemons, A. and Kelley, M. W. (2019). GSK3 regulates hair cell fate in the developing mammalian cochlea. Dev Biol 453, 191–205.

9. Estrach, S., Ambler, C. A., Lo Celso, C., Hozumi, K. and Watt, F. M. (2006). Jagged 1 is a beta-catenin target gene required for ectopic hair follicle formation in adult epidermis. Development 133, 4427–4438.

10. Farin, H. F., Jordens, I., Mosa, M. H., Basak, O., Korving, J., Tauriello, D. V., de Punder, K., Angers, S., Peters, P. J., Maurice, M. M., et al. (2016). Visualization of a short-range Wnt gradient in the intestinal stem-cell niche. Nature 530, 340–343.

11. Hayashi, T., Cunningham, D. and Bermingham-McDonogh, O. (2007). Loss of Fgfr3 leads to excess hair cell development in the mouse organ of Corti. Dev Dyn 236, 525–533.

12. Holley, M., Rhodes, C., Kneebone, A., Herde, M. K., Fleming, M. and Steel, K. P. (2010). Emx2 and early hair cell development in the mouse inner ear. Dev Biol 340, 547–556.

13. Huh, S. H., Jones, J., Warchol, M. E. and Ornitz, D. M. (2012). Differentiation of the lateral compartment of the cochlea requires a temporally restricted FGF20 signal. PLoS Biol 10, e1001231.

14. Hwang, C. H., Guo, D., Harris, M. A., Howard, O., Mishina, Y., Gan, L., Harris, S. E. and Wu, D. K. (2010). Role of bone morphogenetic proteins on cochlear hair cell formation: analyses of Noggin and Bmp2 mutant mice. Dev Dyn 239, 505–513.

15. Jacques, B. E., Puligilla, C., Weichert, R. M., Ferrer-Vaquer, A., Hadjantonakis, A. K., Kelley, M. W. and Dabdoub, A. (2012). A dual function for canonical Wnt/beta-catenin signaling in the developing mammalian cochlea. Development 139, 4395–4404.

16. Jan, T. A., Chai, R., Sayyid, Z. N., van Amerongen, R., Xia, A., Wang, T., Sinkkonen, S. T., Zeng, Y. A., Levin, J. R., Heller, S., et al. (2013). Tympanic border cells are Wnt-responsive and can act as progenitors for postnatal mouse cochlear cells. Development 140, 1196–1206.

17. Jansson, L., Ebeid, M., Shen, J. W., Mokhtari, T. E., Quiruz, L. A., Ornitz, D. M., Huh, S. H. and Cheng, A. G. (2019). beta-Catenin is required for radial cell patterning and identity in the developing mouse cochlea. Proc Natl Acad Sci U S A 116, 21054–21060.

18. Kiernan, A. E., Ahituv, N., Fuchs, H., Balling, R., Avraham, K. B., Steel, K. P. and Hrabe de Angelis, M. (2001). The Notch ligand Jagged1 is required for inner ear sensory development. Proc Natl Acad Sci U S A 98, 3873–3878.

19. Kiernan, A. E., Xu, J. and Gridley, T. (2006). The Notch ligand JAG1 is required for sensory progenitor development in the mammalian inner ear. PLoS Genet 2, e4.

20. Liu, J., Xiao, Q., Xiao, J., Niu, C., Li, Y., Zhang, X., Zhou, Z., Shu, G. and Yin, G. (2022a). Wnt/beta-catenin signalling: function, biological mechanisms, and therapeutic opportunities. Signal Transduct Target Ther 7, 3.

21. Liu, M., Du, Q., Mao, G., Dai, N. and Zhang, F. (2022b). MYB proto-oncogene like 2 promotes hepatocellular carcinoma growth and glycolysis via binding to the Optic atrophy 3 promoter and activating its expression. Bioengineered 13, 5344–5356.

22. McLean, W. J., Yin, X., Lu, L., Lenz, D. R., McLean, D., Langer, R., Karp, J. M. and Edge, A. S. B. (2017). Clonal Expansion of Lgr5-Positive Cells from Mammalian Cochlea and High-Purity Generation of Sensory Hair Cells. Cell Rep 18, 1917–1929.

23. Munnamalai, V. and Fekete, D. M. (2013). Wnt signaling during cochlear development. Semin Cell Dev Biol 24, 480–489.

24. Munnamalai, V. and Fekete, D. M. (2016). Notch-Wnt-Bmp crosstalk regulates radial patterning in the mouse cochlea in a spatiotemporal manner. Development 143, 4003–4015.

25. Munnamalai, V. and Fekete, D. M. (2020). The acquisition of positional information across the radial axis of the cochlea. Dev Dyn 249, 281–297.

26. Munnamalai, V., Hayashi, T. and Bermingham-McDonogh, O. (2012). Notch prosensory effects in the Mammalian cochlea are partially mediated by Fgf20. J Neurosci 32, 12876–12884.

27. Munnamalai, V., Sienknecht, U. J., Duncan, R. K., Scott, M. K., Thawani, A., Fantetti, K. N., Atallah, N. M., Biesemeier, D. J., Song, K. H., Luethy, K., et al. (2017). Wnt9a Can Influence Cell Fates and Neural Connectivity across the Radial Axis of the Developing Cochlea. J Neurosci 37, 8975–8988.

28. Musa, J., Aynaud, M. M., Mirabeau, O., Delattre, O. and Grunewald, T. G. (2017). MYBL2 (B-Myb): a central regulator of cell proliferation, cell survival and differentiation involved in tumorigenesis. Cell Death Dis 8, e2895.

29. Ng, L. F., Kaur, P., Bunnag, N., Suresh, J., Sung, I. C. H., Tan, Q. H., Gruber, J. and Tolwinski, N. S. (2019). WNT Signaling in Disease. Cells 8.

30. Ohyama, T., Basch, M. L., Mishina, Y., Lyons, K. M., Segil, N. and Groves, A. K. (2010). BMP signaling is necessary for patterning the sensory and nonsensory regions of the developing mammalian cochlea. J Neurosci 30, 15044–15051.

31. Oliver, B. L., Young, C. A. and Munnamalai, V. (2021). Spatial and temporal expression of PORCN is highly dynamic in the developing mouse cochlea. Gene Expr Patterns, 119214.

32. Papetti, M. and Augenlicht, L. H. (2011). MYBL2, a link between proliferation and differentiation in maturing colon epithelial cells. J Cell Physiol 226, 785–791.

33. Petrovic, J., Galvez, H., Neves, J., Abello, G. and Giraldez, F. (2015). Differential regulation of Hes/Hey genes during inner ear development. Dev Neurobiol 75, 703–720.

34. Proffitt, K. D. and Virshup, D. M. (2012). Precise regulation of porcupine activity is required for physiological Wnt signaling. J Biol Chem 287, 34167–34178.

35. Routledge, D. and Scholpp, S. (2019). Mechanisms of intercellular Wnt transport. Development 146.

36. Samarajeewa, A., Jacques, B. E. and Dabdoub, A. (2019). Therapeutic Potential of Wnt and Notch Signaling and Epigenetic Regulation in Mammalian Sensory Hair Cell Regeneration. Mol Ther 27, 904-911.

37. Samarajeewa, A., Lenz, D. R., Xie, L., Chiang, H., Kirchner, R., Mulvaney, J. F., Edge, A. S. B. and Dabdoub, A. (2018). Transcriptional response to Wnt activation regulates the regenerative capacity of the mammalian cochlea. Development 145.

38. Shi, F., Cheng, Y. F., Wang, X. L. and Edge, A. S. (2010). Beta-catenin up-regulates Atoh1 expression in neural progenitor cells by interaction with an Atoh1 3’ enhancer. J Biol Chem 285, 392–400.

39. Shi, F., Hu, L., Jacques, B. E., Mulvaney, J. F., Dabdoub, A. and Edge, A. S. (2014). beta-Catenin is required for hair-cell differentiation in the cochlea. J Neurosci 34, 6470–6479.

40. Stevens, C. B., Davies, A. L., Battista, S., Lewis, J. H. and Fekete, D. M. (2003). Forced activation of Wnt signaling alters morphogenesis and sensory organ identity in the chicken inner ear. Dev Biol 261, 149–164.

41. Thompson, M. J., Young, C. A., Munnamalai, V. and Umulis, D. M. (2023). Early radial positional information in the cochlea is optimized by a precise linear BMP gradient and enhanced by SOX2. Sci Rep 13, 8567.

42. Ward, C., Volpe, G., Cauchy, P., Ptasinska, A., Almaghrabi, R., Blakemore, D., Nafria, M., Kestner, D., Frampton, J., Murphy, G., et al. (2018). Fine-Tuning Mybl2 Is Required for Proper Mesenchymal-to-Epithelial Transition during Somatic Reprogramming. Cell Rep 24, 1496–1511 e1498.

43. Whitlon, D. S. (1993). E-cadherin in the mature and developing organ of Corti of the mouse. J Neurocytol 22, 1030–1038.

44. Zak, M. and Daudet, N. (2021). A gradient of Wnt activity positions the neurosensory domains of the inner ear. Elife 10.

