## Supplements for "Sensory progenitors influence patterning of the mammalian auditory sensory epithelium"

**Supplemental Figures:**

**S1:**

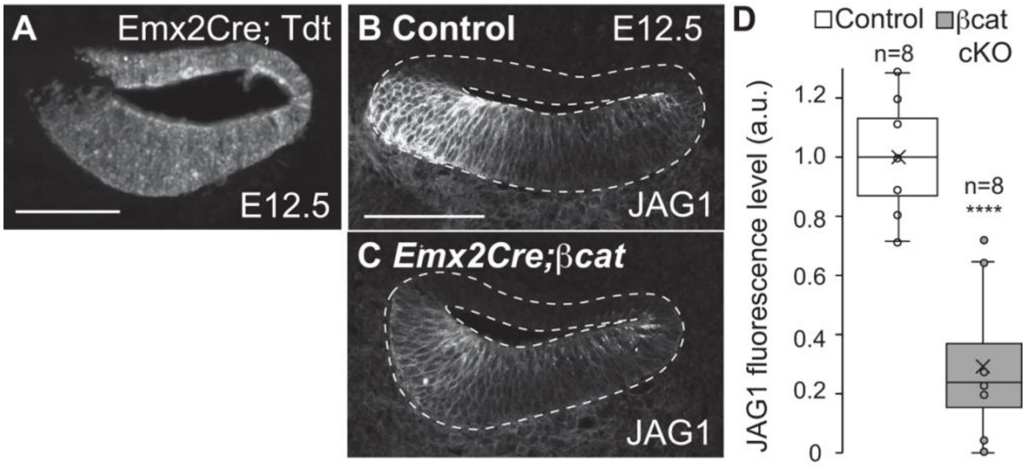


**Supplemental Figure 1: JAG1 is regulated by the Wnt signaling pathway.**

(A) TDT expression upon *Emx2Cre* mediated recombination on E12.5. (B, C) Control cochlea shows JAG1 expression on the medial edge, while early loss of *β-cat* resulted in a decrease in JAG1 expression on E12.5. (D) Quantification of total JAG1 intensity in control and *Emx2Cre*; *β-cat* cKO cochleas on E12.5. (Control N = 8 cochlea, n = 12 sections, *β-cat* cKO N = 8 cochleas, n = 12 sections, Students two-tailed t-test p-value = 1.04e-7). Scale bar = 100μm

**S2:**

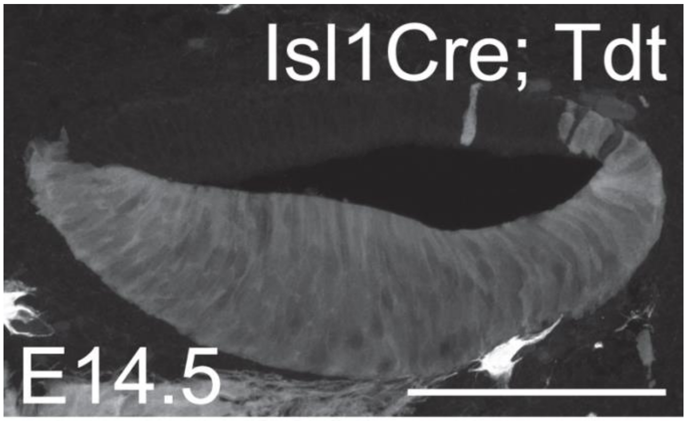


**Supplemental Figure 2: *Isl1Cre* mediated recombination on E14.5.**

TDT expression in the mid turn of the E14.5 cochlea in *Is1lCre*$^{(+/-)}$*;Tdt* mice. Scale bar = 100μm
